# No evidence for the radiation time lag model after whole genome duplications in Teleostei

**DOI:** 10.1101/099663

**Authors:** Sacha Laurent, Nicolas Salamin, Marc Robinson-Rechavi

## Abstract

The short and long term effects of polyploidization on the evolutionary fate of lineages is still unclear despite much interest. First recognized in land plants, it has become clear that polyploidization is widespread in eukaryotes, notably at the origin of vertebrates and teleost fishes. Many hypotheses have been proposed to link the evolutionary success of lineages and whole genome duplications. For instance, the radiation time lag model suggests that paleopolyploidy would favour the apparition of key innovations, although the evolutionary success would not become apparent until a later dispersion event. Some results indicate that this model may be observed during land plant evolution. In this work, we test predictions of the radiation time lag model using both fossil data and molecular phylogenies in ancient and more recent teleost whole genome duplications. We fail to find any evidence of delayed evolutionary success after any of these events and conclude that paleopolyploidization still remains to be unambiguously linked to evolutionary success in fishes.

## Introduction

The understanding of how biodiversity changes and is maintained on Earth has long fascinated biologists. Studying historical biodiversity trends was originally performed by observing the fossil record in successive geological layers [1] and using models to explain the apparition of new clades [2]. New ways of studying the evolution of lineages through time are now available. These involve the joint use of molecular clocks and fossil calibrations in a maximum likelihood [3] or Bayesian [4,5] framework, on the one hand, and of methods enabling evolutionary inferences based on molecular phylogenies [6,7], on the other hand. As the development of complex methodologies retracing evolutionary trends using fossils has also seen continued development [8–10], researchers can now use data from molecules, fossils, or a mix of both when retracing biodiversity [11–13].

Among intrinsic properties of lineages, polyploidy has been the object of intense scrutiny for its potential effect upon evolutionary success of lineages [14–19]. Various scenarios have been proposed to explain the effects of polyploidy on diversification rates. One of them is the radiation time lag model, in which the increase of diversification would occur only after a substantial amount of evolutionary time [20]. The expansion of the lineages would only occur after the evolution of a key innovative trait, thanks to the duplicated gene material, and a subsequent dispersion event. Similarly, Dodsworth et al. [21] have argued that the diploidization process is responsible for the lag between polyploidization and radiation of the lineages. Another scenario, in which the increase in diversification is mediated by reciprocal gene loss leading to reproductive isolation [22], would predict that diversification is highest shortly after the polyploidy event, once differential fixation of paralogs has started in the subpopulations resulting from the polyploid ancestor. Diversification is then predicted to slowly decay as the process of reproductive isolation is complete between all subpopulations. Some evidence for this model had been found in yeast [23]. However, recent modelling work indicates that reciprocal gene loss is not likely to be sufficient for explaining potential diversity increases [24].

In plants, recent findings tend to support the radiation time lag model when considering nine paleopolyploidization events [25]. Using a phylogenetic tree of land plants resolved at the family level, Tank et al. [25] estimated diversification rates and changes using species richness associated with each family. They identified increases in diversification rates in ancestors of the current plant groups and reported that these increases were preferentially clustered after paleopolyploidization. Nevertheless, the estimations of diversification rates were based on family richness, which has been shown to yield a very high rate of false positives [26]. Moreover, it does not model explicitly the changes in diversification rates as a function of time, but rather the global rate at which descendants of a particular internal node of the phylogenetic tree diversify. Finally, our knowledge of plant evolution is rapidly changing with the on-going discovery and dating of paleopolyploidy event in land plants [27–30], and the relation between diversification rates and polyploid will need to be updated accordingly.

Whole genome duplications have also been identified in vertebrates, in particular in the ancestor of all present-day teleost species [31]. Moreover, subsequent polyploidization has also been found in several teleost genera [32], as well as in non-teleost actinopterygian species, such as sturgeons [33]. The Salmoniformes-specific genome duplication has recently been thoroughly studied [34, 35] and dated to occur at least 88 million years ago [36]. Some authors have tentatively linked this genome duplication with anadromy — living in a marine environment and migrating to freshwater to mate — in some salmonid species [37]. Other events of more recent polyploidization have also been investigated in the Cyprinidae family based on the genome of *Cyprinus carpio* [38] or in *Squalius alburnoides* [39], in Botiidae (clown loaches and allies) [40] and in Callichthyidae [41], among others.

Although the occurrence of the ancient genome duplication in actinopterygians coincides with the ancestor to all teleost fishes [42,43], Santini et al. [44] determined, using the same methods as Tank et al. [25], that the teleost-specific genome duplication was responsible for only part of the present-day diversity of this group. By reconstructing diversification patterns, Zhan et al. [45] studied the differences between polyploids and diploids in four groups of actinopterygians and concluded that the impact of polyploidy was inconsistent across these clades. They report a positive impact on cyprinids evolutionary success, but not in salmonids, Botiidae or sturgeons. Similarly, Macqueen et al. [36] report a chronological decoupling between the whole genome duplication of salmonids and their diversification, although without detailing diversification patterns. Moreover, Robertson et al. [46] suggest a delay in re-diploidization affecting approximately one quarter of the genome, following the salmonid-specific whole genome duplication. This would have led to divergence between many ohnologs only after the major salmonid lineages had arisen. These results hint at evolution following the radiation time lag model of diversification in Salmoniformes.

In this work, we propose to further test the assumption of the radiation time lag scenario by studying the responses in diversification in Teleostei after whole genome duplications, comparing them to their sister clades. Our expectation is that we should see a consistent pattern with a surge in diversification sometime after the whole genome duplication event. For this we explicitly model the changes in rates of diversification through time, using both molecular and fossil based knowledge extracted from the literature, on ancient and more recent whole genome duplications.

## Materials and Methods

### Phylogeny-based diversification analysis

We used the data generated from the study of Zhan et al. [45] available at the dryad repository [47]. They reconstructed the phylogenies of four actinopterygian clades: Acipenseridae (sturgeons), Botiidae, Cyprinidae (carp, goldfish and allies) and the monophyletic group formed by the sister orders Salmoniformes and Esociformes (pike and allies). Paleopolyploidization has been identified in the lineage leading to the ancestor Salmoniformes species whereas Esociformes are diploids [34,36]. Similarly, Botiidae is a freshwater family belonging to the Cypriniformes order originating from Southeast Asia with two subfamilies, of which Botiinae is all tetraploid [40]. Sturgeons are also a well-known case of a non-Teleostei group where polyploidy has been recurrently identified [33].

To identify the potential effects of radiation time lag, we compare diversification rates through time between polyploid clades and their sister diploid clades. The identification of changes in diversification through time requires a sufficient number of data points to make meaningful inferences. We thus discarded for our analysis Acipenseridae and 7 events from Cyprinids, because they involved either diploid or polyploid clades of less than 5 species. From the set of polyploidy events under consideration, 5 for cyprinids, 1 for salmonids and 1 for Botiidae, we extracted each polyploid clade and its sister diploid clade. We reconstructed the diversification pattern for each clade independently. We used two different diversification methods, which allow macro-evolutionary rate fitting as a function of time, to compare the dynamics of diversification and highlight potential signs of delayed rise after the paleopolyploidy.

The first method that we used was TreePar [48], which can estimate diversification rates across time ranges on a phylogenetic tree. We have previously shown that this method is robust to mass extinctions, as have affected actinopterygian evolution over the time period considered [49]. The method is run sequentially, starting from a model where only one speciation and extinction rates govern the entirety of the tree. Then, phylogenies are split across a range of time points and macro-evolutionary rates are estimated for each time slice created. The best estimates and time breaks are chosen by their associated log likelihood values. To determine whether a model with an additional break was preferred over the simpler model, we used a likelihood ratio test at *p* ≤ 0.01 significance threshold, according to standard procedure for this method [48]. If the likelihood ratio test favours the simpler model, the computation is stopped and this diversification pattern will be accepted for this tree. If the more complex model is preferred, the computation is run once more with one additional time break, until a model is accepted. Using this procedure thus enables us to find abrupt changes in diversification at certain time points.

The second method used was developed by Morlon et al. [50]. It enables fitting any function to speciation and extinction rates using time as the variable. We tested three different responses that assumed either constant, linear or exponential relationship between rates and time. We fit each type of function for each rate, for a total of 9 models, and chose the one with the best fit to our data using ΔAIC, as employed by Morlon et al. [50]. This method will be referred to as the function-fitting method.

We ran both analyses on the distribution of 500 trees that we extracted from the dryad repository [47] for each group of species, to check whether all phylogenetic trees agreed onto a similar diversification scenario. We performed the diversification analyses on the full distribution of trees of Botiidae and salmonids species, and on five subclades of cyprinids showing differential ploidy level between sister clades. All scripts used for this part of the analysis are available at https://github.com/cashalow/teleost-radiation.

### Fossil-based approach

We downloaded from the Paleobiology database (https://paleobiodb.org/, accessed Feb. 2016) all actinopterygian fossil occurrences identified at the species level, separating teleosts and non-teleosts. For non-teleosts, we selected occurrences matched to the Chondrostei, Cladistia or Holostei taxonomic groups. We obtained 1,239 and 1,515 occurrences for species from teleosts and non-teleosts, respectively. Each species name present in our datasets that matched accepted names from Fish-Base [51] was assumed to be extant, while the others were considered extinct.

We used PyRates (https://github.com/dsilvestro/PyRate) [8] to estimate diversification rates from fossil occurrences of extinct and extant species without prior knowledge of relationships between the considered species. Additionally to speciation and extinction, preservation and sampling rates for the lineages are also estimated, so that the differences of likelihood of being observed in the fossil record between species do not bias the macro-evolutionary rate estimations. On top of this probabilistic framework, a Bayesian procedure is used to explore models with different numbers of changes in macro-evolutionary rates. Thus, from the total distribution of the dating of the occurrences, we are able to reconstruct the changes in diversification rates through geological time, with both extinct and extant species fossils.

## Results

### Salmonids, cyprinids and botiids whole genome duplications studied with phylogeny methods

Overall, there is no signal for a radiation time-lag, although in some clades there are differences in diversification rates between polyploids and diploids. Let us first consider the case where diversification was determined to be constant inside both diploid and polyploid clades (figure 1 and figure 2 panels B, C, D and E). In most cases, both methods are consistent, whether supporting higher diversification rates in polyploids (figure 2 panels B,C and D) or no difference (figure 2 panel E). The exception is Salmonids, for which TreePar strongly supports that polyploids diversified faster, whereas function-fitting provides overlapping ranges of estimated diversification values. Of note, the subtree noted ‘E’ in figure 2 represents the youngest whole genome duplication event studied in this work, and no difference is found between polyploids and diploids. We note that in some cases, individual trees led to incoherent inferences relative to the majority inference for the same species, when using the function-fitting method: 1 tree in figure 1 and 1 tree in figure 2 (D) were fit with exponentially decaying instead of constant diversification functions.

**Fig 1.**
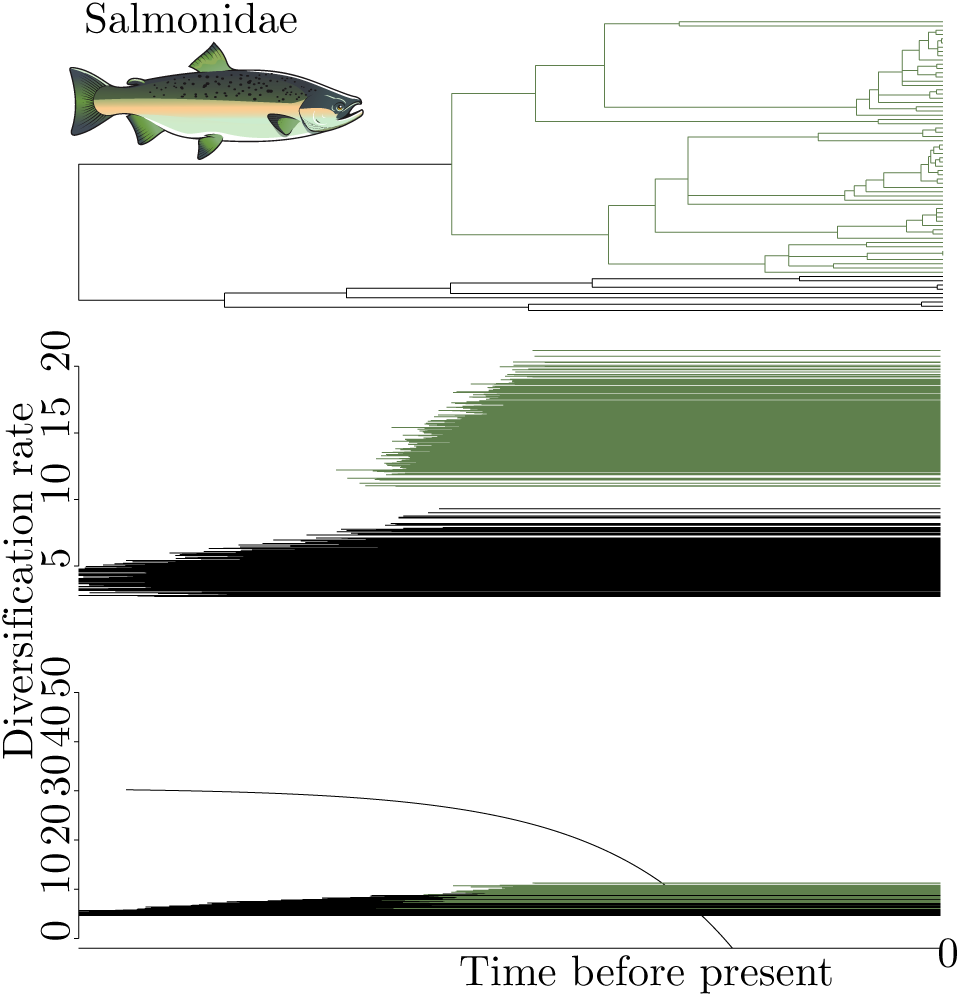
Results for the Salmoniformes and Esociformes phylogeny. The black clade represents the diploid Esociformes, the dark green clade the tetraploid Salmoniformes. Top: consensus phylogenetic tree; middle: TreePar analysis; bottom: function-fitting. One line represents the result for one diploid or polyploid clade extracted from one the 500 hundred trees. Constant diversification models were preferred with every analysis, as no TreePar results includes a break in diversification value, and as only constant functions were chosen by the function-fitting method except for one outlier diploid clade. The analysis were run with the sampling ratios extracted from Zhan et al. [45]: 0.69 for the polyploid clade and 0.28 for the diploid one.

**Fig 2.**
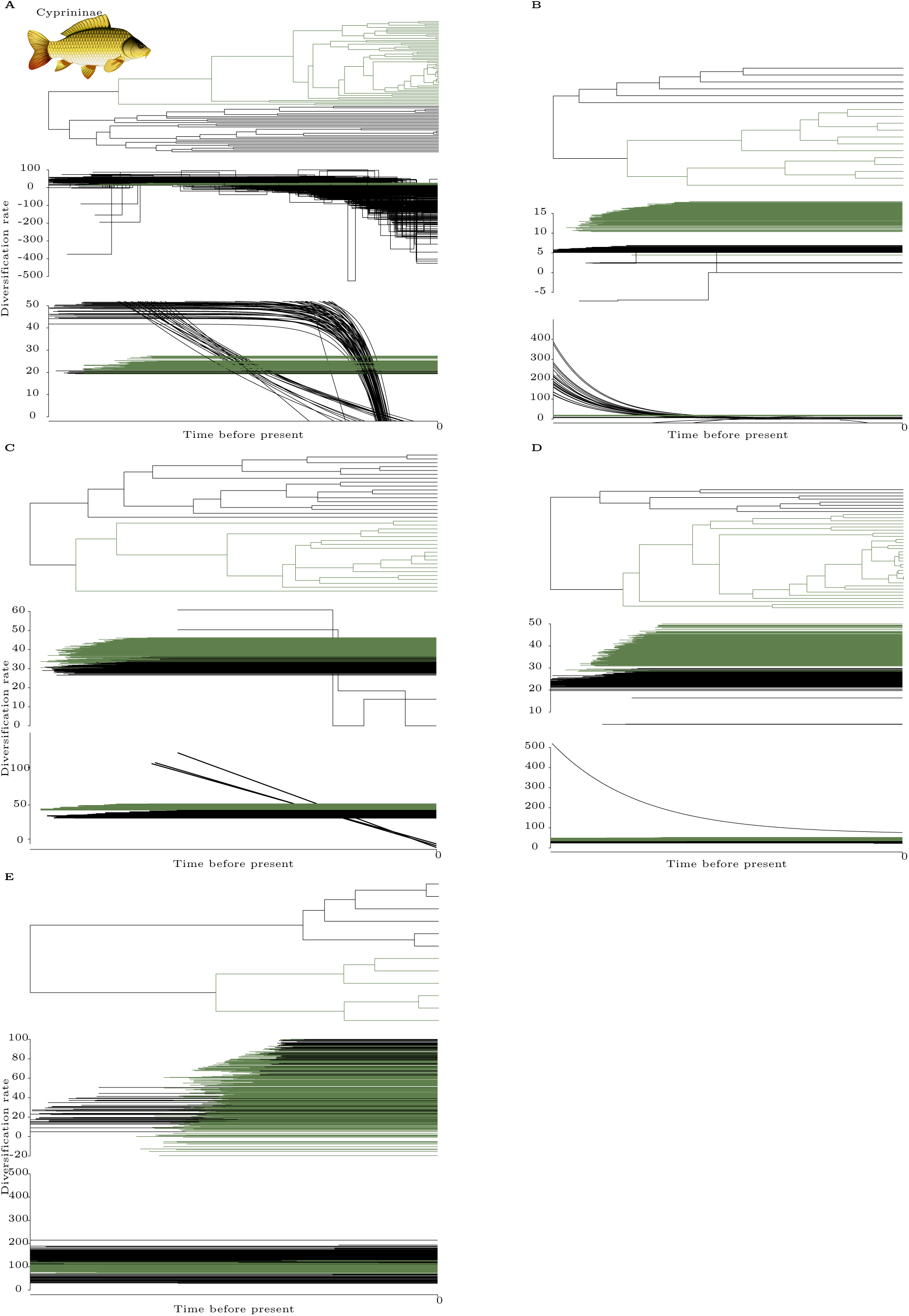
(Next page) Results for five Cyprininae subtrees, ordered from oldest to most recent event, presented as in figure 1. No fossil calibration was performed thus the absolute timing of the events cannot be estimated. Most genera were recovered as paraphyletic in [45], except *Capoeta, Pseudobarbus, Schizothorax* and *Sinocyclocheilus,* genera for which monophyly had been documented before. Lack of monophyly prevents the estimation of the sampling ratio of each subtrees, thus the analysis were run without sampling information. Panel A: results comparing a diploid clade (black), encompassing *Gara* and *Labeo* genera, with a tetraploid one (dark green) encompassing the *Barbus, Labeobarbus, Neolissochilus, Varicorhinus* and *Tor* genera. Panel B: subtrees encompassing *Pseudobarbus* and *Barbus* species. Panel C: subtrees with hexaploid *Capoeta* genus members and tetraploid *Barbus* and *Luciobarbus* species. Panel D: results from a subtree of panel A, comprising *Casobarbus* genera and some *Barbus* species in the tetraploid subclade (black), and hexaploid species of *Labeobarbus* and *Barbus* genera (dark green). Panel E: comparison inside the *Schizothorax* genus.

When diversification was inferred to be changing through time, consistent scenarios were reconstructed by both methods (figures 3 and 2 panel A). For the oldest duplication in cyprinids, the best model for TreePar assumed a constant diversification rate for polyploid, while diploids diversification rates decrease from initially faster than polyploids to slower. With function-fitting, a similar scenario is observed for most of the trees of the distribution, with some variability for diploids. Indeed, this method favours the following scenarios for diploids: most trees support an exponential decrease in diversification, either (i) gradually or (ii) more sharply and closer to the present; a small proportion support (iii) constant diversification, at a lower rate than the polyploids. In all cases, polyploids are consistently diversifying faster than diploids near the present. For Botiidae, the pattern is similar. Diploids initially diversified faster and then reached the approximate levels of diversification of polyploids. For TreePar, this was modelled for the vast majority of trees as a two-phase diversification value for the diploid clade only, with a switch in rates near the present at roughly the same time for every tree, and a constant diversification rate for polyploids. For function-fitting, this was modelled as an exponential decay for most of the distribution of the diploid clade, and a constant value of diversification the remainder. The exponential decay led to lower rates of diploids than polyploids close to the present, whereas the constant values has higher rates for diploids. Like for constant rate phylogenies, we also observe rare inconsistencies for a few trees for both methods.

**Fig 3.**
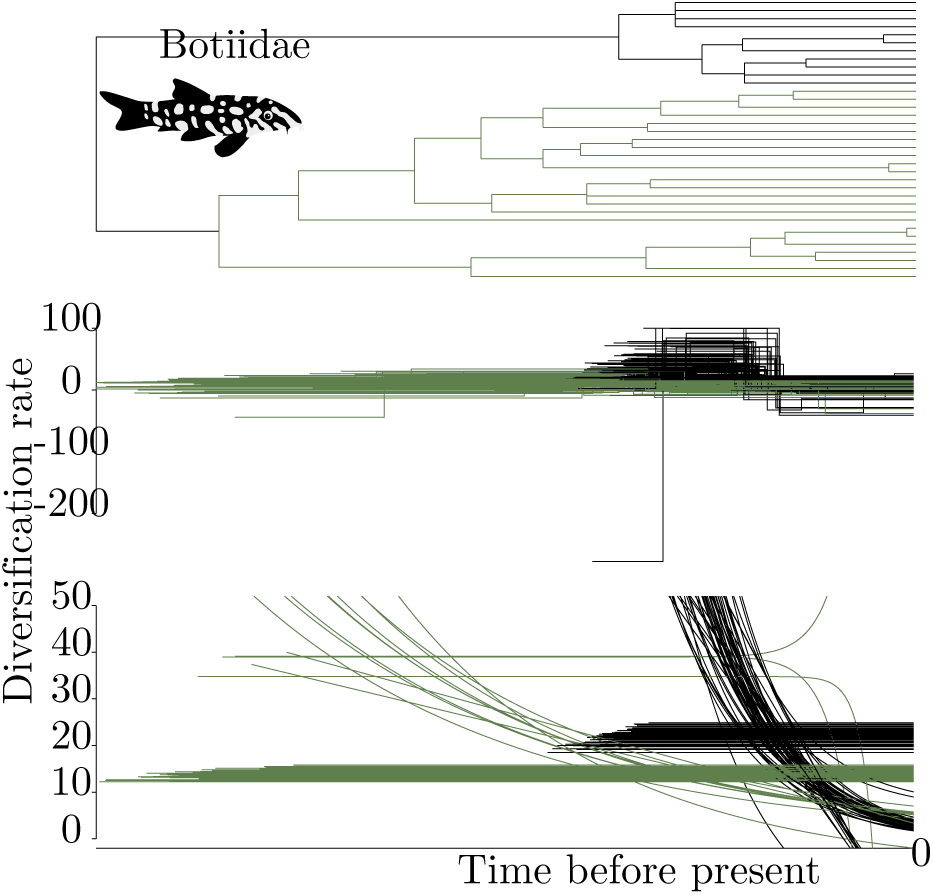
Results for the Botiidae (clown loaches and allies) species, presented as in figure 1.

### Teleost whole genome duplication studied with fossil data

From the origin of non-teleost actinopterygians till the Permian-Triassic extinction event, the diversification rate for these species stayed constant and at a relatively high level (figure 4, black line), level that were never reached again in subsequent 250 My of evolution. Their diversification rate then plunged below zero, around the Permian-Triassic mass extinction event (250 Mya), denoting massive loss of species, and then stabilized over the long run near null diversification, explaining the relative rarity of those species in the present day.

**Fig 4.**
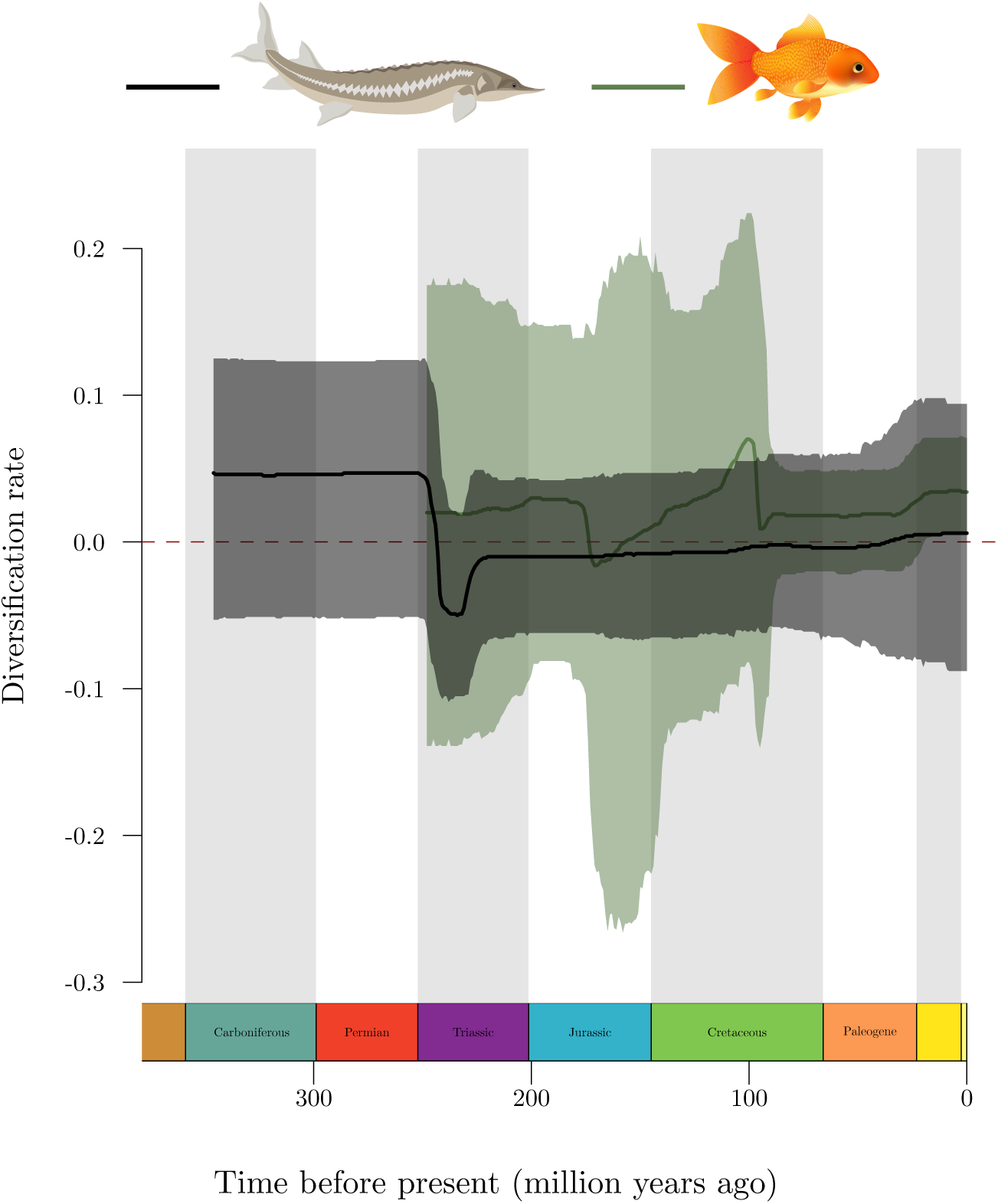
Results of the fossil analysis, for species-matched fossils. In black is the reconstructed diversification mean (thick line) and 95% highest density interval (transparent area) from fossil occurrences of the Chondrostei, Cladistia, or Holostei groups, including as extant species bichirs, bowfins, gars or sturgeons (pictured) and allies, which did not experience paleopolyploidy. In dark green is the results of the same analysis for the fossil attributed to Teleostei, encompassing most of the approximate 30000 extant species, among them cyprinids and goldfishes (pictured), whose last ancestor experienced polyploidization.

Although teleost fishes appeared around the boundary of the Carboniferous and Permian periods (298 Mya) [52], we are not able to estimate diversification rates before 250 Mya because of lack of data. From 250 Mya, teleost fishes diversified at a constant rate for more than 50 Mya, faster than their non-teleostean counterparts around the same time but at lower levels than the latter experienced before Teleostei appeared, during the Carboniferous and Permian periods. They experienced a sudden drop in their diversification in the middle of the Jurassic (around 175 Mya), consistent with the reported loss of 53 genera of ray finned fishes around this boundary [53]. They went on to a steady increase in their diversification during the Early Cretaceous but around 100 Mya, their diversification once again sharply decreased, possibly driven by marine specific groups [53].

Interestingly, most of the ray finned fish families disappearing at the Cretaceous-Paleogene boundary were exclusively marine, whereas no fully freshwater family disappeared [53]. After this last drop, Teleostei diversification rate stayed constant at a positive value until the present. The confidence intervals on all these estimates are very large, such that it is difficult to conclude on the significance of variations, over time and between teleosts and non-teleosts.

## Discussion

In this study, we tested the radiation time lag model after whole genome duplications in teleost fishes. For this model to be verified, an increase in diversification rate should be seen, and it should be some time after the whole genome duplication rather than at the time of the event. We have thus compared both recent and older events, used both fossil and phylogenetic data, and compared the fates of polyploid clades to their sister diploid clades. We used methods with different initial assumptions in order to test their consistency across datasets. We did not find evidence for the radiation time lag model in any case that we investigated.

Both methods were mostly consistent in the patterns of diversification that they supported. We note that although our study does not support the radiation time-lag model, some clades appear to follow a somewhat related scenario for diversification. Cyprinids support a pattern where diploid diversification is initially higher than polyploids but then decays until it reaches lower values. The radiation time lag model posits that key innovations occur because of the whole genome duplication, but that the radiation and the appearance of new species happen only once the lineage carrying the key innovation is dispersed to some new environment [20]. This hypothesis should lead to an increase in diversification for polyploids, not a decrease in diploids. Nevertheless, these results show a potential case where the evolutionary advantage of polyploids over diploids is not apparent before a significant amount of time. In that way, it is similar to the radiation time lag model.

Paleopolyploidization events in cyprinids or Botiidae have not been precisely dated, and it is difficult to estimate whether sufficient time has elapsed for the postulated effects of the radiation time lag to be observed. Nevertheless, in every cyprinid and Botiidae cases that we investigated, constant diversification was observed after the polyploidy event. Even though in some cases this polyploid event led to higher diversification compared to diploids (figure 2 panel B, C and D), this pattern does not support the radiation time lag model because the increase in diversification appears concomitantly with the polyploidization.

The second oldest event investigated concerns the salmonids. Contrary to what Macqueen et al. [36] reported when studying the diversification of Salmoniformes, we did not find increasing diversification rates with time, but a constant rate of diversification for both polyploid salmonids and diploids Esociformes, using both TreePar and the function-fitting method. The salmonid specific genome duplication has been dated between 88 Mya [36] and 96 Mya (90.5-101.5) [34], whereas the common ancestor to every extant Salmonidae species is usually dated around 55 Mya (52.2-58.0 [52], 52.1-59.5 [54]). Based on a lineage through time analysis, Macqueen et al. [36] concluded that the pattern of diversification was more consistent with an increased diversification correlated to the cooling down of the ocean. But since they did not estimate diversification rates, their results cannot support that diversification increased through time. Overall, we do not find evidence of delayed increase of diversification after the salmonid specific duplication.

The importance of the anadromous behaviour in salmonids has already been reported [36] and tentatively linked with the whole genome duplication [37]. Macqueen et al. [36] performed a BiSSE analysis on the phylogeny of the Salmoniformes, discriminating diversification rates between anadromous and fully freshwater species, to test for a correlation between evolutionary success and this behaviour. They only reported the estimation of the speciation rate of anadromous and freshwater species, and although anadromous species had higher speciation, higher extinction in these species could still lead to a non-significant or deleterious effect upon diversification. Alexandrou et al. [37] found that anadromy evolved multiple times, and no sooner than 55 My after the whole genome duplication. That pattern might fit the prediction of the radiation time lag, if the evolution of such behaviour would lead to increase in diversification. Nevertheless, in our analysis, we found no differences in diversification rates even after the appearance of anadromy, and with one of the two method used, we even found similar diversification rates between Salmoniformes and Esociformes (figure 1, bottom panel), highlighting the lack of evidence for such a mode of diversification.

Most of the results found here are in partial agreement with previous studies [45], where differences were investigated by studying the trees as a whole and discriminating between ploidy levels using BiSSE [55], rather than separately analysing each clade by ploidy-level. Indeed, their results also indicated higher speciation rates of salmonids. Nevertheless, they reported higher diversification for Botiidae polyploids, whereas we do not find significant difference between the two clades. Comparisons between the results are complicated by the fact that the methods used in this study allow to model more complex evolutionary scenarios than BiSSE does. There is also some debate about the statistical properties of the BiSSE method [56], which warrants caution when interpreting results obtained by this method.

The teleost whole genome duplication has been alternatively dated around 350 Mya [57], between 300 and 450 Mya [58], or between 226 and 316 Mya [59]. The date of the most recent common ancestor to all teleosts was also recently re-evaluated, with estimates ranging from 307 Mya (285-333) [52], to 283 Mya (255-305) [60]. There is thus a minimum lag of 50 My between the occurrence of the genome duplication and the appearance of the common ancestor of all extant teleosts. In our results, the first estimations of diversification are available from around 250 Mya, meaning that fossil occurrences are not frequent enough between this date and the whole genome duplication to estimate diversification rates.

In the oldest event in our dataset, common to all teleosts, there was no increase in diversification found before at least 50 My after the appearance of the first teleost fossil, and only after a sharp decrease in diversification. Moreover, the diversification rate of teleosts went back up to the initial value more than 100 My later, and never quite reached the rates at which non-Teleostei species were diversifying across the Permian and Carboniferous. This seems to indicate that the tremendous diversity observed today in Teleostei is the result of a steady diversification process over geological time, rather than an increased diversification promoted by the whole genome duplication.

The effect of the teleost whole genome duplication had previously been studied by looking a the fossil record [19], but without explicit modelling of the processes of origination of families, by studying the probability of survival of families emerging before and after whole genome duplications, and hypothesized that whole genome duplication acted as a protection against extinction of such families. In our study, we explicitly modelled the diversification rates and found no signal indicating a protection against extinction after whole genome duplication over the long run in teleost fishes. Recently, the evolution of new organs, such as the bulbus arteriosus which is specific to the teleost heart, has been linked to the whole genome duplication [61]. Both sub— and neofunctionalization of a ohnolog duplicate elastin gene is thought to have enabled the apparition of the bulbus arteriosus. Although this organ had tentatively been linked to the success of Teleostei, we fail to find such a signal in their diversification rate.

Overall, through the 8 different events studied here, none showed support for the radiation time lag model; 3 events supported higher diversification in polyploids (all belonging to Cyprininae) and one showed inconsistent results between the two methods (Salmonidae).

## Acknowledgments

This work was supported by the ProDoc grant number 134931 of the Swiss National Science Fondation; and État de Vaud. The computations were performed at the Vital-IT (http://www.vital-it.ch) Center for high-performance computing of the Swiss Institute of Bioinformatics. We would like to thank Christian Parisod and Nils Arrigo for comments on an earlier version of the manuscript.

